# Asymmetric Turnover Dynamics of Nav1.6 Voltage-Gated Sodium Channels at the Axon Initial Segment

**DOI:** 10.64898/2026.09.10.747750

**Authors:** Daisuke Yoshioka, Kohei Yamamoto, Lei Huang, Manabu Abe, Kenji Sakimura, Takafumi Kawai, Yoshifumi Okochi, Yasushi Okamura

## Abstract

Nav1.6 voltage-gated sodium channels are critical for shaping action potentials, and their precise localization at the axon initial segment (AIS) is essential for neuronal excitability. However, the turnover mechanisms that maintain this spatial pattern remain unclear. Here, we present a genetically engineered mouse in which endogenous Nav1.6 carries a Cre-switchable fluorescent tag, enabling us to simultaneously trace the turnover of pre-existing and newly synthesized Nav1.6 without perturbing AIS structure. Using this system, we determine the physiological lifetimes of Nav1.6 at the AIS in vivo and in vitro, and further uncover the spatially asymmetric turnover dynamics, with the proximal and distal AIS organizing a source-to-sink gradient. Based on the quantified parameters, computational modeling clarifies that this asymmetry promotes efficient clearance of older molecules, thereby supporting AIS quality control. Collectively, these findings provide a technical and conceptual foundation for understanding how molecular turnover contributes to AIS homeostasis.

## Main

The axon initial segment (AIS) plays a crucial role in establishing and maintaining neuronal polarity, and serves as the central hub for the initiation and modulation of action potentials (AP) (1–3). Activity-dependent AIS plasticity further supports neuronal homeostasis by tuning output to changes in excitability (4–7). Membrane depolarization and AP firing are driven by voltage-gated sodium (Nav) channels (8). Direct visualization of Na^+^ flux and computational modeling have demonstrated that a high density of functional sodium channels at the AIS is required for AP generation (9–11). Among the nine Nav channel paralogs encoded in the mammalian genome (Nav1.1–Nav1.9), four subtypes (Nav1.1, Nav1.2, Nav1.3, and Nav1.6) are abundantly expressed in the brain and exhibit distinct cell-type specificity and subcellular distributions (12,13). Notably, Nav1.6 is the predominant subtype universally localized to the AIS of mature neurons across multiple neuronal populations (14–19). The significance of Nav1.6 in AP regulation is further underscored by the diverse neurological phenotypes associated with its loss or mutation, including juvenile lethality, paralysis, ataxia, tremor, muscle weakness, and dystonia (20).

To regulate appropriate excitability in long-lived neurons, the AIS structure must be organized in accordance with intrinsic activity and external stimuli. The spatial distribution of AIS proteins such as Nav1.6 is defined by the balance among three-dimensional trafficking processes, including exocytosis, lateral diffusion, and endocytosis (21). Accordingly, modulation of the molecular turnover constitutes a fundamental mechanistic basis of AIS plasticity (4–7). Even at steady state, the finite lifetimes of molecules make their dynamic and continuous exchange essential for preserving the structural and functional integrity of the AIS. Although recent advances in live imaging of endogenous AIS proteins have been remarkable (6,22,23), current knowledge of AIS organization and dynamics still largely derives from bulk measurements of molecular density, leaving unresolved how Nav1.6 turnover proceeds within the AIS and how it supports long-term neuronal homeostasis. This knowledge gap stems from technical limitations of conventional approaches. Stable isotope labeling combined with mass spectrometry enables comprehensive quantification of protein lifetimes at the whole-brain level (24–26), but lacks subcellular spatial resolution. Short hairpin RNA (shRNA)-based knockdown strategies can conveniently assess decay kinetics across diverse target proteins (27), yet suppression of protein expression itself perturbs AIS structure, precluding accurate measurement of physiological protein lifetimes at the AIS. Moreover, none of these methods can capture the appearance process of newly synthesized molecules. Although short-term imaging has clearly visualized preferential exocytosis of Nav1.6 at the AIS followed by rapid diffusion confinement (28,29), these observations do not explain the long-term turnover dynamics required to regulate AIS integrity. Substantial gaps therefore remain in our understanding of how Nav1.6 turnover is organized within the AIS to support neuronal homeostasis.

Here, we aim to elucidate the molecular mechanisms that govern the formation and maintenance of Nav1.6 distribution at the AIS. To this end, we generated a genetically engineered mouse line (Nav1.6-FLEx mice) in which endogenous Nav1.6 carries a Cre-switchable fluorescent tag, enabling non-perturbative visualization of Nav1.6 replacement history. Using this approach, we quantitatively determined the time constants that define physiological Nav1.6 turnover in vivo and in vitro, revealing a previously unrecognized source– sink organization in which molecular appearance and disappearance preferentially occur at the proximal and distal AIS regions. Furthermore, computational modeling based on these measurements clarifies that the spatially asymmetric turnover enables a density-gradient-driven clearance mechanism that efficiently removes older molecules, thereby contributing to AIS quality control. These findings establish a technical and conceptual foundation for an integrated understanding of how Nav1.6 turnover is spatiotemporally organized to maintain AIS homeostasis.

## Results

### Cre-switchable tagging enables non-perturbative visualization of Nav1.6 turnover at the AIS

To directly visualize the spatiotemporal dynamics of Nav1.6 turnover without perturbing AIS, we first established a genetically engineered mouse line (Nav1.6-FLEx mice) in which the fluorescent tag on endogenous Nav1.6 can be switched from EGFP to tdTomato in a Cre-dependent manner via a Flip-excision (FLEx) system (Fig. 1a). In this system, the EGFP signal marks older channels that already existed before Cre expression, whereas the tdTomato signal marks those newly synthesized afterward. Using confocal imaging, we confirmed the Cre-dependent switch from Nav1.6-EGFP to Nav1.6-tdTomato expression in hippocampal neurons of Nav1.6-FLEx mice in vivo and in vitro following infection with adeno-associated virus (AAV) carrying calmodulin-dependent protein kinase II (CaMKII)-Cre (AAV-CaMKII-Cre) (Fig. 1b). The recombinant Nav1.6 colocalized with ankyrinG (ankG), a master organizer of AIS, and was shifted distally relative to ankG (Fig. 1c–e), as previously reported (17,18,28). These observations indicate that recombinant Nav1.6 normally localizes to the AIS.

**Fig. 1:**
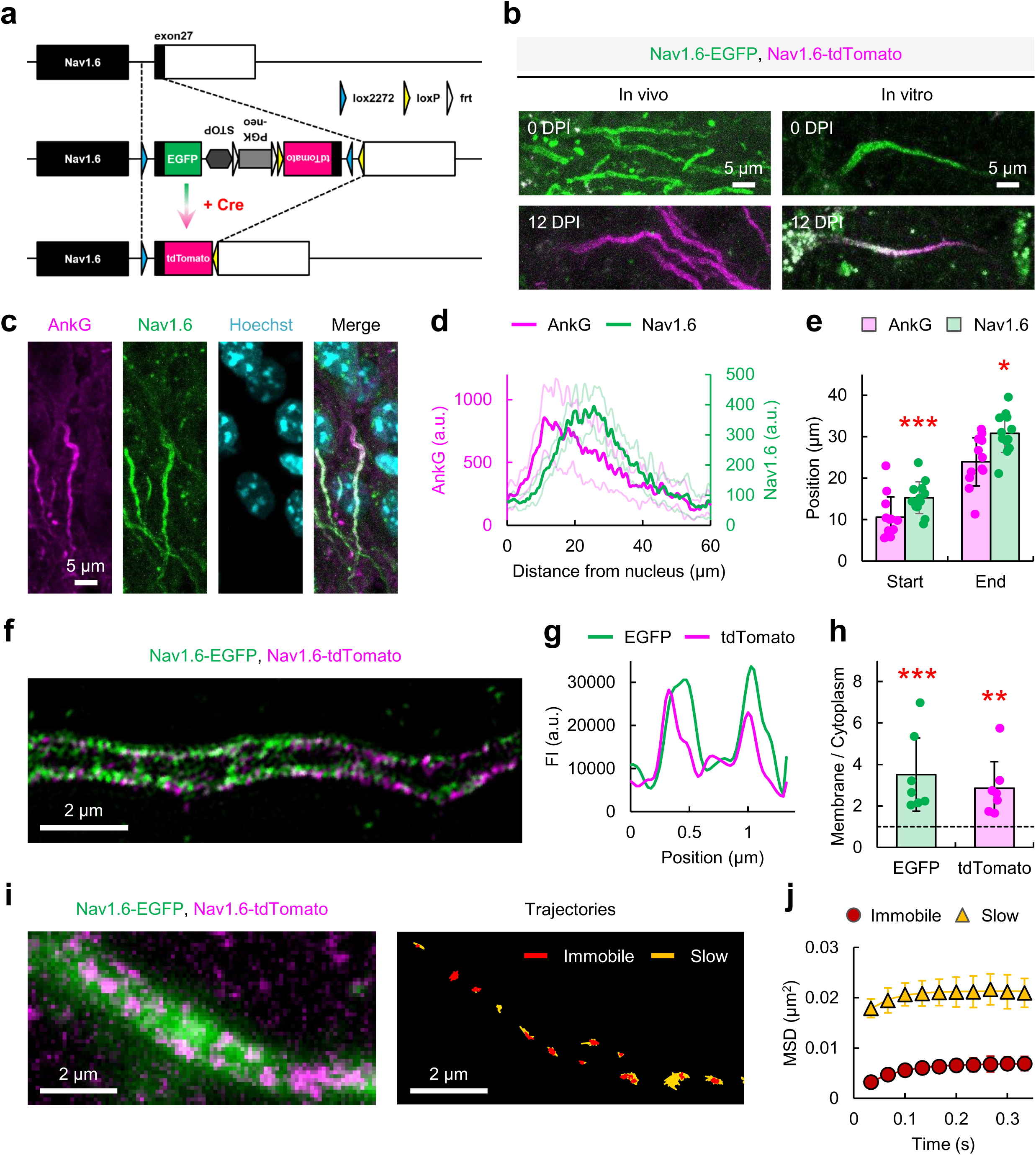
Cre-switchable tagging enables non-perturbative visualization of Nav1.6 turnover while preserving its normal AIS distribution and diffusion dynamics. **a,** Schematic of the Cre-dependent switch of the fluorescent tag on Nav1.6 using the FLEx system. **b**, Representative confocal images of fixed hippocampal sections and primary cultured hippocampal neurons expressing Nav1.6-EGFP or Nav1.6-tdTomato at 0 and 12 DPI. Scale bar, 5 μm. **c**, Representative confocal images of Nav1.6-EGFP-expressing neurons in fixed hippocampal sections immunostained for ankG. Nuclei were marked by Hoechst. Scale bar, 5 μm. **d**,**e**, Fluorescence intensity profiles and the start (\*\*\**P* = 0.001) and end (\**P* = 0.012) positions of ankG and Nav1.6-EGFP along the axon, measured from the edge of the nucleus (*n* = 12). *P*-values were obtained by paired *t* test. **f**, Representative SIM image of the AIS in primary cultured hippocampal neurons at 4 DPI (21 DIV) expressing Nav1.6-EGFP and Nav1.6-tdTomato. Scale bar, 2 μm. **g**, Representative fluorescence intensity profiles of Nav1.6-EGFP and Nav1.6-tdTomato along the short axis of the axon. **h**, Fluorescence intensity ratio between the plasma membrane and the cytoplasm (*n* = 7). EGFP, \*\*\**P* = 0.000; tdTomato, \*\**P* = 0.002 (paired *t* test for membrane versus cytoplasm). **i**, Representative EPI image of Nav1.6-EGFP and TIRF image of Nav1.6-tdTomato expressed in a primary cultured hippocampal neuron at 4 DPI (21 DIV). The diffusion trajectories for 40 s of Nav1.6-tdTomato are shown. Red and yellow lines indicate trajectories of immobile and slow-diffusion states of the molecules within an interval of 33 ms, respectively. Scale bar, 2 μm. **j**, Ensemble MSD versus time plots for immobile and slow-diffusion states of Nav1.6-tdTomato at the AIS of primary cultured hippocampal neurons at 4 DPI (21 DIV) (*n* = 5). **d** shows mean ± 95% CI; **e**, **h**, and **j** show mean ± SD.

Structured illumination microscopy (SIM) further confirmed that Nav1.6-EGFP and Nav1.6-tdTomato localize predominantly to the plasma membrane rather than the cytoplasm (Fig. 1f–h). Thus, the AIS fluorescence signals in confocal images primarily reflect the surface density of Nav1.6 molecules. To examine the lateral diffusion dynamics of recombinant Nav1.6 at the AIS, we performed single-molecule imaging using total internal reflection fluorescence microscopy (TIRFM). In neurons at 4 d post-infection (DPI), when Nav1.6-tdTomato expression was still low, we detected sparsely distributed fluorescent spots corresponding to newly inserted Nav1.6-tdTomato molecules or clusters (Fig. 1i, Left). Their trajectories were classified into two diffusion states (immobile and slow) based on the Akaike information criterion (AIC) and a hidden Markov model (HMM) (Fig. 1i, Right; see Methods) (30,31). Mean squared displacement (MSD) analysis (Eq. 1) showed that Nav1.6 diffusion was confined in both states (Fig. 1j and Extended Data Fig. 1), consistent with previous reports (28,29). These results demonstrate that recombinant Nav1.6 preserves normal surface expression and nanoscale dynamics.

We also evaluated the channel functionality of recombinant Nav1.6. Because the channel function of Nav1.6-EGFP has already been reported to be indistinguishable from that of the wild-type channel (32), we focused here on characterizing the channel properties of Nav1.6-tdTomato. Whole-cell patch-clamp recordings revealed no significant differences in current density or in gating properties for activation and inactivation between non-tagged and tdTomato-tagged Nav1.6 (Extended Data Fig. 2). Combined with the previous report (32), these results indicate that neither EGFP nor tdTomato affects the channel properties of Nav1.6. Taken together, our data demonstrate that recombinant channels in Nav1.6-FLEx mice retain normal spatial patterns, diffusion dynamics, and channel properties. Therefore, this approach enables visualization of the homeostatic replacement of older channels with newly synthesized channels without perturbing AIS structure.

### Non-perturbative visualization of Nav1.6 turnover reveals its physiological lifetime at the AIS

Having established that recombinant Nav1.6 is functionally localized to the AIS, we proceeded to quantify its turnover. In our in vivo experiments, we defined a measurement window between P61 and P97 during which the morphological features of Nav1.6 distribution remained stable, indicating an apparent steady state (Extended Data Fig. 3a). After AAV-CaMKII-Cre infection, we acquired confocal images of fluorescence intensity profiles of Nav1.6-EGFP and Nav1.6-tdTomato along the axon at 0, 6, and 12 DPI (Fig. 2a). This approach enabled us to simultaneously visualize the decay of pre-existing Nav1.6-EGFP and the accumulation of newly synthesized Nav1.6-tdTomato within the same AIS following Cre expression. The decay kinetics of EGFP signals across the entire Nav1.6 distribution were fitted with a single-exponential function, yielding a time constant (**τ**) of 12.3 d (Fig. 2b). Because AAV infection does not necessarily occur simultaneously across neurons and a delay exists before the consequent tag switch, the estimated **τ**should be regarded as a conservative upper bound on the true value.

**Fig. 2:**
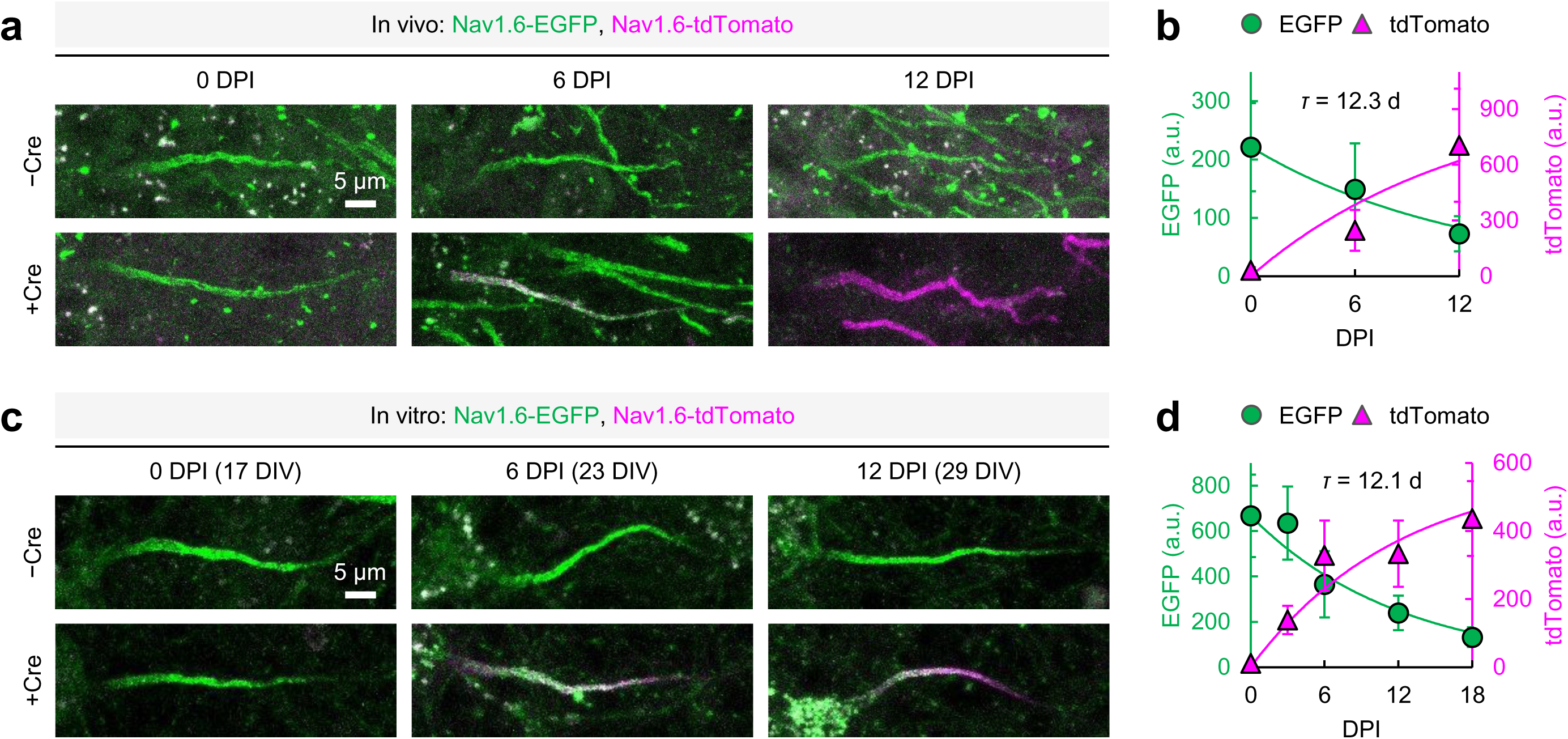
Non-perturbative visualization of Nav1.6 turnover reveals its physiological lifetime at the AIS. **a**, Representative confocal images of neurons expressing Nav1.6-EGFP or Nav1.6-tdTomato in fixed hippocampal sections at 0, 6, and 12 DPI. Scale bar, 5 μm. **b**, Mean fluorescence intensities of Nav1.6-EGFP and Nav1.6-tdTomato along the axon (10–50 μm from the edge of the nucleus) in fixed hippocampal sections at 0, 6, and 12 DPI (*n* = 11–29). The Nav1.6-EGFP data were fitted with a single-exponential function (**τ**= 12.3 d). The Nav1.6-tdTomato data were fitted with an inverse single-exponential function, with **τ**fixed to the value obtained from the Nav1.6-EGFP data and only *F*_max_ estimated (*F*_max_ = 995.15 a.u.). **c**, Representative confocal images of primary cultured hippocampal neurons expressing Nav1.6-EGFP or Nav1.6-tdTomato at 0, 6, and 12 DPI. **d**, Mean fluorescence intensities of Nav1.6-EGFP and Nav1.6-tdTomato along the axon (0–40 μm from the edge of the soma) in primary cultured hippocampal neurons at various DPI (*n* = 17–33). The Nav1.6-EGFP data were fitted with a single-exponential function (**τ**= 12.1 d). The Nav1.6-tdTomato data were fitted with an inverse single-exponential function, with **τ**fixed to the value obtained from the Nav1.6-EGFP data and only *F*_max_ estimated (*F*_max_ = 591.95 a.u.). Data are mean ± SD.

In our in vitro experiments, AAV-CaMKII-Cre was applied at 17 DIV, when Nav1.6 distribution was fully established (Extended Data Fig. 3b). Consistent with the in vivo results, we again observed exponential decay and accumulation of Nav1.6 signals across the entire distribution, yielding a **τ**of 12.1 d (Fig. 2c,d). These findings indicate that the characteristic turnover rate of Nav1.6 is largely independent of external factors such as myelin sheaths or extracellular matrix, instead reflecting intrinsic neuronal mechanisms. Furthermore, in both in vivo and in vitro systems, the **τ**obtained from the decay kinetics of EGFP signals also described the accumulation kinetics of tdTomato signals (Fig. 2b,d). This consistency supports the experimental validity that the sequential process of AAV infection, Cre expression, and tag switch did not differentially perturb the turnover of Nav1.6-EGFP and Nav1.6-tdTomato. Collectively, our approach enables quantitative determination of the effective lifetime of Nav1.6 at the AIS under physiological conditions without perturbing AIS structure.

### Spatially asymmetric Nav1.6 turnover forms a source-to-sink gradient along the AIS

Building on our determination of the overall turnover time constant, we next characterized the spatiotemporal dynamics of Nav1.6 turnover in more detail. We first analyzed the time-dependent changes in the morphological properties of Nav1.6 distribution. In vivo, the Nav1.6-EGFP distribution progressively shortened from the distal end (33.459 ± 5.983 μm) over time, whereas the proximal end (15.832 ± 5.153 μm) remained unchanged (Fig. 3a,b). This pattern indicates that Nav1.6 disappearance occurs preferentially at the distal AIS. In contrast, during the initial rising phase that primarily reflects molecular accumulation, the centroid of the Nav1.6-tdTomato distribution (23.197 ± 5.449 μm) was aligned with that of the entire Nav1.6 distribution as defined by the 0-DPI EGFP signal (24.770 ± 5.089 μm), being proximally shifted from the distal end where disappearance is prominent (Fig. 3a–c). Thus, this pattern indicates that Nav1.6 appearance is promoted preferentially at the proximal AIS relative to its disappearance. The spatially asymmetric turnover dynamics of Nav1.6, characterized by distal disappearance coupled with proximal appearance, were robustly reproduced in vitro as well (Fig. 3d–f).

**Fig. 3:**
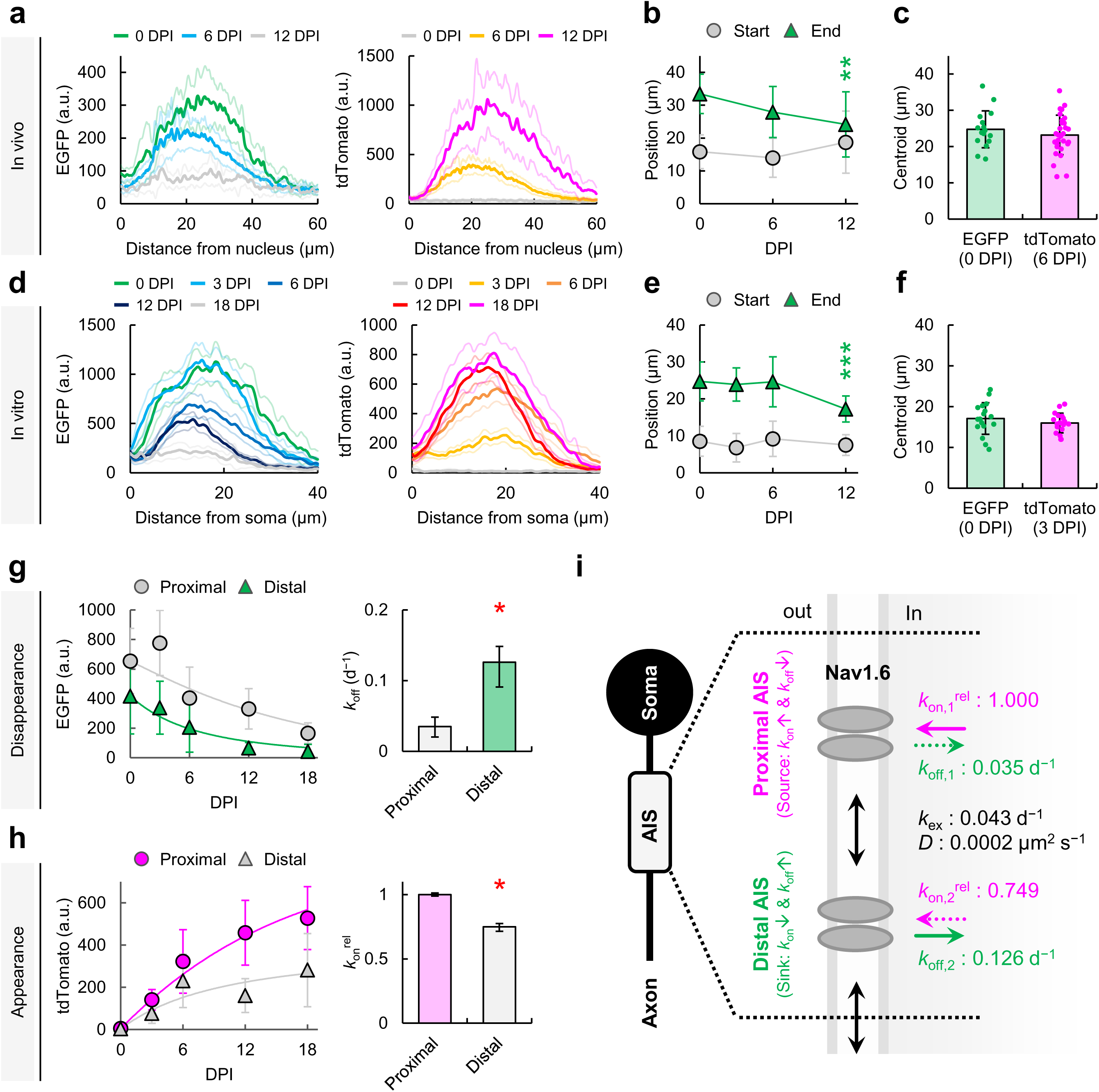
Spatially asymmetric Nav1.6 turnover forms a source-to-sink gradient along the AIS. **a,b,** Fluorescence intensity profiles and the start and end positions of the Nav1.6-EGFP distribution along the axon, measured from the edge of the nucleus, in fixed hippocampal sections with Cre expression at 0, 6, and 12 DPI (*n* = 11–29). \*\**P* < 0.01 (Dunnett’s test, comparisons were made with the “0 DPI” data). **c**, Centroids of the Nav1.6-EGFP distribution at 0 DPI (*n* = 16) and the Nav1.6-tdTomato distribution at 6 DPI (*n* = 29). *P* = 0.353 (Welch’s *t* test). **d**,**e**, Fluorescence intensity profiles and the start and end positions of Nav1.6-EGFP along the axon, measured from the edge of the soma, in primary cultured hippocampal neurons with Cre expression at various DPI (*n* = 17–33). \*\**P* < 0.01 (Dunnett’s test, comparisons were made with the “0 DPI” data). **f**, Centroids of the Nav1.6-EGFP distribution at 0 DPI (*n* = 17) and the Nav1.6-tdTomato distribution at 3 DPI (*n* = 18). *P* = 0.327 (Welch’s *t* test). **g**,**h**, Mean fluorescence intensities of Nav1.6-EGFP and Nav1.6-tdTomato in the proximal (0–20 μm from the soma) and distal (20–40 μm from the soma) regions of the AIS in primary cultured hippocampal neurons at various DPI (*n* = 17–33). Disappearance and appearance curves were fitted with Eqs. 2, 3, 6, and 7 to quantify *k*_off_ and *k*_on_^rel^ at the proximal and distal AIS. \**P* < 0.05 (the 95% BCa CI for the difference between proximal and distal parameters did not include zero). **a** and **d** show mean ± 95% CI; **b**, **c**, **e**, and **f** show mean ± SD; disappearance and appearance curves in **g** and **h** also show mean ± SD; and *k*_off_ and *k*_on_^rel^ in **g** and **h** are presented with 95% BCa CI. **i**, Schematic models summarizing the quantified parameters underlying asymmetric turnover dynamics of Nav1.6 at the proximal and distal AIS.

We next sought to quantify the rate constants that govern the asymmetric Nav1.6 turnover. For this analysis, we used in vitro data, which provided greater experimental stability and therefore higher statistical reliability than in vivo data. Based on the centroid of the Nav1.6 distribution (17.087 ± 3.863 μm; Fig. 3f), the AIS was divided into proximal (0–20 μm) and distal (20–40 μm) regions. The decay time course of EGFP signals in each region was fitted using a compartmental kinetic model (Eqs. 2 and 3; see Methods), yielding the disappearance rate constant (*k*_off_) and the exchange rate constant (*k*_ex_) (Fig. 3g). Because Nav1.6 molecules predominantly reside on the axonal surface (Fig. 1f–h), *k*_off_ reflects the efficiency of molecular disappearance through endocytic internalization. The estimated *k*_off_ was significantly higher in the distal region (0.126 d^−1^, 95% BCa CI 0.091–0.148 d^−1^) than in the proximal region (0.035 d^−1^, 95% BCa CI 0.020–0.048 d^−1^), indicating preferential disappearance at the distal AIS (Fig. 3g,i). *k*_ex_ represents the rate constant governing molecular exchange between the proximal and distal compartments, accounting for the lateral diffusion of Nav1.6 along the axon (Fig. 1i,j and Extended Data Fig. 1) (28,29). Following previous reports showing no detectable shuttle of Nav1.6 or other AIS channels (e.g., Kv2.1) along the cell surface between the soma and axon (28,33), we set no molecular exchange at the proximal end of AIS in this model. Using Fick’s first law (Eqs. 4 and 5; see Methods), we estimated the diffusion coefficient (*D*) from *k*_ex_ (0.043 d^−1^, 95% BCa CI 0.019–0.059 d^−1^) to be 0.0002 μm^2^ s^−1^, 95% BCa CI 8.9×10^−5^–2.7×10^−4^ μm^2^ s^−1^ (Fig. 3i), comparable to previously reported values (0.0007 ± 0.0006 μm^2^ s^−1^, *n* = 20) (28).

Finally, with *k*_off_ and *k*_ex_ fixed at the values obtained from the decay kinetics of the full dataset, we quantified the apparent appearance rate constant (*k*_on_^app^) by fitting the accumulation time course of tdTomato signals in each region using a compartmental kinetic model (Eqs. 6 and 7; see Methods). As Nav1.6 is mainly present on the axonal surface (Fig. 1f–h), *k*_on_^app^ reflects the efficiency of molecular appearance through exocytic insertion. The relative appearance rate constant (*k*_on_^rel^), normalized to *k*_on_^app^ at the proximal region from the full dataset, was significantly lower in the distal region (0.749, 95% BCa CI 0.715–0.775) than in the proximal region (1.000, 95% BCa CI 0.988–1.013), indicating preferential appearance at the proximal AIS (Fig. 3h,i). Supporting this conclusion, in early-stage immature neurons where Nav1.6 expression is not yet prominent (14,15,17,28), Nav1.6 begins to accumulate from the proximal AIS, and its distribution progressively extends toward the distal region over time (Extended Data Fig. 4). In summary, these findings demonstrate that the spatial distribution of Nav1.6 is established and maintained by spatially asymmetric turnover dynamics in mature neurons under basal conditions. Specifically, Nav1.6 accumulates more actively at the proximal AIS through exocytic insertion, whereas its removal through endocytic internalization is more active at the distal AIS (Fig. 3i). Thus, within a single AIS, the proximal and distal regions asymmetrically function to form a source-to-sink gradient.

### Asymmetric turnover dynamics preferentially drive older Nav1.6 from the proximal to the distal AIS

What is the physiological significance of the asymmetric turnover dynamics described above? Lateral diffusion of AIS channels between the soma and axon is extremely rare or virtually absent (28,33), suggesting that a slow, unidirectional flux following the molecular density gradient could arise along the axon from the proximal toward the distal side. This raises the hypothesis that molecules drift preferentially from the proximal to the distal AIS depending on their age on the axonal surface.

To examine how asymmetric turnover shapes the age composition of molecular populations within the proximal and distal AIS, we performed single-molecule Monte Carlo simulations parameterized by experimentally derived kinetic constants (Fig. 3i; see Methods). The simulations reproduced both the temporal evolution of molecule numbers at the proximal and distal AIS and the spatial density gradient observed at 120 d, a time point approximating steady state (Fig. 4a,b). These results support the validity of the estimated parameters and the simulation model. We then defined the effective molecular age as the time between molecular appearance and disappearance at the AIS and applied a demographic framework to assess the contribution of each age group to population formation in distinct AIS subregions. To visualize the relationship of molecular displacement and age, we classified molecules present at steady state (120 d) according to their region of origin and generated heatmaps of the joint probability for molecular position and age, P(x, age) (Fig. 4c). Molecules exiting and entering the proximal AIS both tended to be older at greater distances from their appearance site, but the absolute number of net migrating molecules was larger for efflux than influx at the proximal AIS. The net migration rate (NMR), defined as the difference between efflux and influx normalized by the total number of molecules in the proximal AIS (Eq. 8), yielded 0.116 ± 0.004 (*n* = 5). Finally, we quantified the age-specific net migration contribution (ANMC), defined as the number of age-specific net migrants normalized by the total number of net migrants (Eq. 9). The ANMC distribution was shifted toward older ages relative to the age distribution of the proximal AIS population P(age | prox) (Fig. 4d). Accordingly, the ANMC-weighted mean age (21.266 ± 0.338 d, *n* = 5), representing the effective age centroid of molecules contributing to net migration, was significantly higher than the mean age of the proximal population (14.462 ± 0.055 d, *n* = 5) (Fig. 4e). Overall, these findings demonstrate that the density-gradient-driven net flux preferentially expels older Nav1.6 molecules from the proximal to the distal AIS (Fig. 4f). This model suggests that the asymmetric turnover may serve as an efficient clearance mechanism for aged Nav1.6, thereby contributing to AIS quality control.

**Fig. 4:**
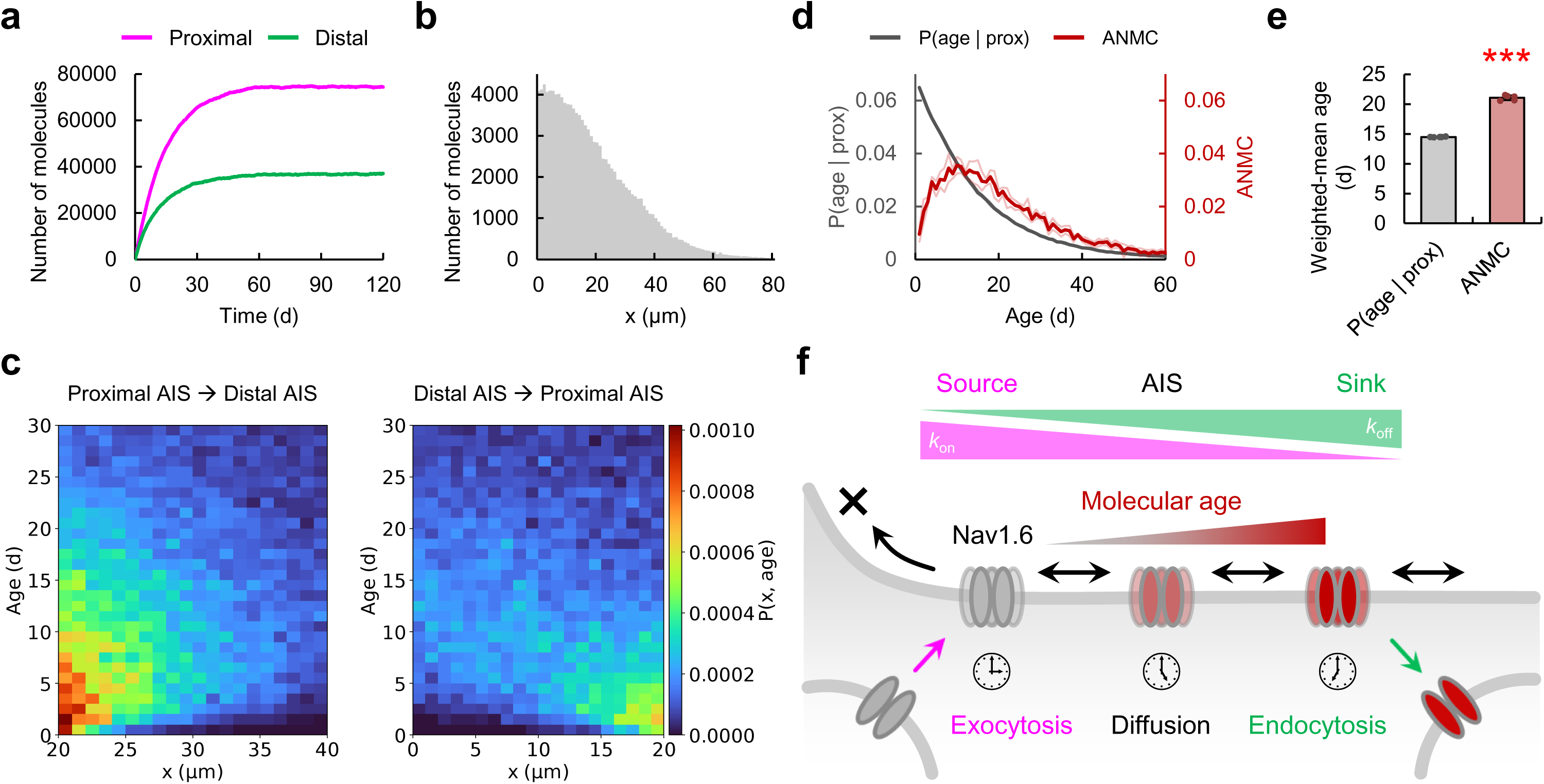
Asymmetric turnover dynamics preferentially drive older molecules from the proximal to the distal AIS. All panels show simulations based on the measured kinetic parameters in each compartment summarized in Fig. 3i. The proximal and distal AIS were defined as x = 0-20 μm and x = 20-40 μm, respectively. **a**, Time courses of the number of molecules in the proximal and distal AIS. **b**, Spatial distribution of molecules along the x-axis at steady state (120 d). **c**, Heatmaps of the joint probability for molecular position and age, P(x, age), computed for molecules originating from the proximal and distal AIS, corresponding to outflow from and inflow to the proximal AIS, respectively. **d**, Probability distribution of molecular age at the proximal AIS, P(age | prox), and ANMC (*n* = 5). **e**, Weighted mean ages of P(age | prox) and ANMC shown in **d**. \*\*\**P* = 0.000 (Welch’s *t* test). Data are mean ± SD. **f**, Schematic of an efficient clearance mechanism for aged Nav1.6. Asymmetric turnover dynamics generate a slow, density-gradient-driven flux from the proximal (source) to the distal (sink) AIS that intrinsically biases migration toward the removal of older molecules. This mechanism may help prevent excessive accumulation of aged Nav1.6, thereby contributing to AIS quality control essential for long-term neuronal homeostasis.

## Discussion

Beyond recent endogenous labeling approaches for AIS proteins that avoid overexpression artifacts (6,22,23), we established a Nav1.6-FLEx system that enables direct visualization of Nav1.6 turnover dynamics without perturbing the AIS (Fig. 1). This system allowed us to quantitatively determine the time constant governing the formation and maintenance of the spatial distribution of Nav1.6 in neurons. The effective mean lifetime of Nav1.6 across its entire distribution was approximately 12 d (half-life ∼8.3 d) (Fig. 2), which is comparable to the previously reported half-life (≥ 14 d) obtained using an shRNA-based approach (27). Our quantitative analytical approach further revealed a spatial asymmetry in Nav1.6 turnover kinetics within a single AIS (Fig. 3). One candidate factor that could underlie this disparity is ankG, a master organizer of AIS architecture (34). AnkG plays a multifaceted role by promoting the exocytic insertion of its binding partners to the AIS (28,31) and by inhibiting their endocytic internalization (31,35). Given that Nav1.6 colocalizes slightly distal to the peak of ankG distribution (Fig. 1c–e) (17,18,28), it is reasonable to attribute enhanced molecular appearance to the ankG-rich proximal region and accelerated disappearance to the ankG-poor distal region. The submembrane periodic skeleton may also contribute to the bidirectional dynamics of Nav1.6 by regulating exo/endocytosis at the AIS (36–38). Nevertheless, the three-dimensional trafficking dynamics of Nav1.6 along the axon remain insufficiently resolved. Thus, future studies will require higher-resolution approaches, such as in-depth single-molecule analyses (31), to more directly probe the trafficking dynamics and mechanisms underlying AIS turnover.

The asymmetric turnover mechanism of Nav1.6 suggests that the functional machineries responsible for its appearance and disappearance are regionally segregated, a configuration that is highly advantageous for preventing physical interference or competition between them. In line with this idea, recent super-resolution imaging has shown that exo-and endocytic sites along the axon are spatiotemporally segregated and rarely occupy the same clearings of the submembrane periodic skeleton (36). Our finding of a proximal bias in the primary source is also mechanistically compatible with the direction of kinesin-mediated anterograde transport of Nav1.6 toward the AIS (39). Furthermore, our computational modeling indicates that asymmetric turnover can function as an efficient clearance mechanism in which a density gradient drives a net flux that selectively expels older molecules from the proximal to the distal AIS (Fig. 4). In general, proteins are exposed to various chemical and physical stresses after their synthesis, leading to the gradual accumulation of molecular damage. For example, reactive oxygen and nitrogen species can alter Nav channel function and thereby affect Na^+^ currents in hippocampal neurons (40). Such molecular aging processes raise the possibility that the asymmetric turnover may contribute to AIS quality control by managing the age distribution of Nav1.6. In future studies, it would be intriguing to explore whether perturbations of turnover mechanisms and the resulting accumulation of aged molecules are associated with neuronal pathology or aging.

The Nav1.6 turnover model established here under basal conditions serves as a robust template for explaining the mechanisms that regulate its distribution across a wide range of steady-state and perturbed conditions. Previous studies have shown that activity-dependent homeostatic plasticity can be observed under various experimental conditions and has been examined in detail (4–7). Structural alterations of the AIS have also been reported in numerous neurological and psychiatric disorders, including Alzheimer’s disease (AD), epilepsy, multiple sclerosis, and bipolar disorder, as well as in neural injuries such as ischemic stroke and optic nerve crush (3). Moreover, AIS remodeling has been observed during mammalian hibernation, a non-pathological process that has been noted to share features with early-stage AD (41). In addition, aging has been reported to shorten the AIS across multiple brain regions (42–44). However, the molecular turnover underlying these various forms of AIS plasticity remains poorly understood. Thus, applying our Nav1.6-FLEx mice to quantify shifts in the balance of Nav1.6 turnover will offer a unified framework for describing how its spatial pattern is established, maintained, and modified across diverse physiological and pathological contexts. Importantly, this methodology can also be directly extended to analyze the turnover of Nav1.6 that accumulates at nodes of Ranvier (45,46), which are essential for AP propagation. Taken together, our study provides a powerful platform to bridge the long-standing gap between phenomenological views of Nav1.6 distribution and a mechanistic insight into its turnover dynamics.

## Methods

### Animals

The use of mice was approved by the Animal Care and Use Committee of The University of Osaka (gene recombination experiment approval number 04636; animal experiment approval number 02-068) and Niigata University (gene recombination experiment approval number SD02311; animal experiment approval number SA01798). Mice were maintained on a 12/12-h light/dark cycle under controlled temperature and humidity, with ad libitum access to food and water. In vitro cultures were prepared from P0–2 mice, and in vivo experiments were performed using P61–97 mice.

### Generation of Nav1.6-FLEx mice

We used a Flip-excision (FLEx) strategy to generate a Cre-dependent fluorescent tag switch at the C-terminus of endogenous Nav1.6 (Fig. 1a). A FLEx cassette was assembled containing two versions of the final exon of *Scn8a* (exon 27): one fused in-frame with a linker (GTRILQSTVGTAGPGSIAT) (32), EGFP, and the human CGF polyadenylation signal, followed by the other fused with the same linker, tdTomato, and the rabbit β-globin polyadenylation sequence (47) in the reverse orientation. A reversed STOP cassette composed of the yeast *His3* terminator and the SV40 polyadenylation signal (48) was inserted downstream of the EGFP-containing exon. A reversed neomycin resistance cassette driven by the phosphoglycerate kinase promoter (PGK-neo) and flanked by flippase recognition target (frt) sites (49) was placed between the reversed STOP cassette and the reversed tdTomato-containing exon. The entire construct was flanked by alternating lox2272 and loxP sites, enabling sequential recombination and excision events that irreversibly flip the cassette in the presence of Cre, thereby switching expression from Nav1.6-EGFP to Nav1.6-tdTomato at the endogenous locus. The 5’ and 3’ homology arms (8.7 kb and 1.5 kb, respectively) were incorporated to facilitate homologous recombination. The PacI-linearized targeting vector was electroporated into RENKA C57BL/6 embryonic stem (ES) cells (50), and G418-resistant clones containing the FLEx cassette were isolated. Correctly targeted ES clones, identified by Southern blot analysis, were injected into eight-cell-stage ICR embryos. Male chimeras were crossed with C57BL/6N females to establish the Nav1.6-FLEx mouse line. The forward (GGATAAGCCTGCTCACTGAG) and reverse (TGAGAGAGGAACAGAGGATG) primers were designed on both sides of the lox2272 sequence for genotyping.

### DNA constructs

The mouse Nav1.6-tdTomato plasmid was generated at VectorBuilder. The same linker used in the Nav1.6-FLEx mouse was positioned between Nav1.6 and tdTomato. In addition, an immunoglobulin (Ig) C ⊥ intron (110 bp) (51) was inserted at position 3140 of the Nav1.6 cDNA to reduce the inhibitory effect of this gene on bacterial growth (52,53). A plasmid encoding non-tagged Nav1.6 was constructed by deleting the linker and tdTomato as follows: The forward primer (GCAGCATCGCCACACTCGAGACCCAGCTTTCTTGTA) was designed to match the sequence containing the XhoI site downstream of tdTomato. The reverse primer (AAAGCTGGGTCTCGAGTGTGGCGATGCTGCCTGGTC) was designed to introduce a stop codon (TAG) and an XhoI site immediately downstream of Nav1.6. The fragment containing Nav1.6 but lacking the linker and tdTomato was amplified by standard PCR using PrimeSTAR Max DNA Polymerase (Takara) and ligated after DpnI and XhoI digestion.

The plasmid for packaging CaMKII-Cre into AAV (pAAV-CaMKII-Cre) was constructed as follows: the CaMKII-Cre fragment was excised from the vector (pENN.AAV.CamKII 0.4.Cre.SV40, Addgene #105558) at the NdeI/HindIII sites and subcloned into the same sites in another vector (YA1531: pAAV_CAG-FLEX-QuasAr3, Addgene #107701), which has ITR. The DNA sequences of all constructs were verified through Sanger sequencing.

### AAV preparation and stereotaxic injections

AAV-CaMKII-Cre was produced in AAVpro 293T cells (Takara) as previously described (54). The cells were cultured in DMEM supplemented with 10% FBS and 0.1% penicillin/streptomycin at 37 °C, 5% CO_2_, and passaged every 2–3 d. For virus production, the cells in 10-cm dishes were transfected with three plasmids (pAAV-CaMKII-Cre; pAAV2/2 rep/cap, Addgene #232199; and pHelper, Takara) using 1 mg mL^−1^ polyethylenimine (PEI; Polysciences Inc.) for 24 h, followed by incubation for an additional 2 d at 35 °C, 3% CO_2_. At 3 d post-transfection, the cells were harvested, lysed by freeze–thaw cycles, and centrifuged. The supernatant was loaded onto iodixanol (OptiPrep, Serumwerk Bernburg AG) density gradients (15%, 25%, 40%, 58%) and fractionated by centrifugation at 16,100 × g for 4 h at 4 °C. The 40% iodixanol fraction containing packaged AAV was collected. Viral titers were determined by quantitative PCR (55), and aliquots were stored at −80 °C until use. The resulting laboratory-produced AAV-CaMKII-Cre (2.1–3.2 × 10^12^ vg mL^−1^) was used for in vivo experiments. In addition, a commercially available AAV1 carrying pENN.AAV.CamKII 0.4.Cre.SV40 (7.2 × 10^12^ vg mL^−1^, Addgene #105558-AAV1) was used for in vitro experiments.

Mice anesthetized with medetomidine (0.75 mg kg^−1^, Sandoz), midazolam (4 mg kg^−1^, Sandoz), and butorphanol (5 mg kg^−1^, Meiji) were secured in a stereotaxic frame (Narishige). 1 μL of AAV solution was injected into the CA2/3 region at coordinates relative to bregma (AP = −1.0 mm, ML = ±1.5 mm, DV = −1.5 mm) using a 26-G syringe (Hamilton). AAV was infused at 0.2 μL min^−1^ for 5 min, followed by an additional 5-min post-injection period. Antisedan (0.75 mg kg^−1^, Zenoaq) was administered intraperitoneally to facilitate rapid recovery from anesthesia.

### Primary culture of hippocampal neurons

Primary hippocampal neuron cultures were prepared as previously described (31). Only the key differences relevant to this study are noted below. Hippocampi were dissected from Nav1.6-FLEx mouse pups (P0–2), and tissues from both male and female pups were pooled. At 17 DIV, neurons were infected with AAV at 1,000–2,000 MOI.

### Confocal imaging

For in vivo experiments, mice were sacrificed and perfused with 1% PFA in PBS. After 1 h of post-fixation in 1% PFA, the brain was transferred to 30% sucrose and incubated overnight at 4 °C. The cryoprotected brain was embedded in O.C.T. compound (Sakura) and frozen at −80 °C. Sections (20 μm thick) were cut in the sagittal plane using a cryomicrotome (Leica Microsystems), mounted onto MAS-coated glass slides (Matsunami), and dried at room temperature for at least 2 h. After washing off the O.C.T. compound with PBS, the sections were incubated with the primary antibody (anti-Ank-G, 1:200, Frontier Institute Co., LTD.) in PBS supplemented with 3% goat serum and 0.2% Triton X-100 overnight at 4 °C. After three washes with PBS (3 min each), the sections were incubated with the secondary antibody (Alexa Fluor 647 anti-rabbit IgG, 1:1,000, Invitrogen) in PBS for 1 h at room temperature. Nuclei were labeled with Hoechst 33258 (1:1,000, Dojindo). After three additional washes with PBS (3 min each), Fluorescence Mounting Medium (Dako) was applied, and the sections were sealed with NEO cover glass (Matsunami).

For in vitro imaging, primary cultured hippocampal neurons were washed three times with 1 mL of artificial cerebrospinal fluid (aCSF: 125 mM NaCl, 5 mM KCl, 1 mM MgCl2, 2 mM CaCl2, 10 mM glucose, 10 mM HEPES, pH 7.4 adjusted with NaOH) and mounted in a magnetic chamber (CM-B18-1, Live Cell Instrument).

Confocal images were acquired using an FV3000 system (Evident) equipped with oil-immersion objectives (UPLSAPO100X, NA 1.45, Evident for in vivo; UPLSAPO60X, NA 1.42, Evident for in vitro). Pixel sizes were 124 nm (in vivo) and 207 nm (in vitro), with a frame size of 1,024 × 1,024 pixels. Imaging was performed at room temperature.

### Analysis of confocal images

Acquired images were processed and analyzed using Fiji/ImageJ (v1.53c). A Gaussian filter (2 pixels) was applied for noise reduction. Images were generated by merging z-stacks with a step size of 0.50 μm for in vivo samples and 0.41 μm for in vitro samples. To quantify the start, end, and length of the Nav1.6-EGFP and Nav1.6-tdTomato distributions, fluorescence intensity profiles along axons were min–max normalized. A threshold of 0.5 was applied, and the start, end, and length of the longest contiguous region above the threshold were extracted. For in vivo data, because the soma was hard to identify under our imaging conditions, fluorescence profiles were plotted from the edge of the nucleus labeled with Hoechst. Mean fluorescence intensities of Nav1.6-EGFP and Nav1.6-tdTomato after background subtraction were obtained from regions defined by the quantified morphological properties of the Nav1.6 distribution (Fig. 3a–c): 10–30 μm (proximal), 30–50 μm (distal), and 0–60 μm (entire) from the nuclear edge. For neurons at P61–97, to exclude clearly immature neurons and reduce artificial variability arising from developmental heterogeneity, only neurons in which the fluorescence signal was continuously sampled without missing values up to 60 μm from the nuclear edge were analyzed. For in vitro data, mean fluorescence intensities of Nav1.6-EGFP and Nav1.6-tdTomato after background subtraction were obtained from regions defined by the quantified morphological properties of the Nav1.6 distribution (Fig. 3d–f): 0–20 μm (proximal), 20–40 μm (distal), and 0–40 μm (entire) from the soma edge. For neurons at 17–35 DIV, to exclude immature neurons and minimize artificial dispersion, only neurons in which the fluorescence signal was continuously sampled without missing values up to 40 μm from the soma edge were analyzed. The decay kinetics of EGFP signals across the entire Nav1.6 distribution were fitted with a single-exponential function, *F*_max_exp(-*t*/**τ**), where *t* is the DPI, **τ** is the time constant, and *F*_max_ is the fluorescence intensity fixed to the value at 0 DPI. The accumulation kinetics of tdTomato signals across the entire Nav1.6 distribution were fitted with an inverse single-exponential function, *F*_max_(1 - exp(-*t*/**τ**)), where *t*, **τ**, and *F*_max_ denote the DPI, the time constant, and the maximum fluorescence intensity, respectively. For this fit, **τ** was fixed to the value obtained from the decay kinetics.

### SIM imaging

Primary cultured hippocampal neurons at 4 DPI (21 DIV) were washed three times with 1 mL of aCSF and fixed with 4% PFA in PBS. The coverslip was then mounted in a magnetic chamber. SIM images were acquired using an ELYRA S1 system (Carl Zeiss) equipped with an oil-immersion objective (alpha Plan-Apochromat 100x/1.46 Oil DIC, 100*, NA 1.46, Carl Zeiss). Pixel sizes were 25 nm, with a frame size of 1,904 * 1,900 pixels. Imaging was performed at room temperature.

Acquired images were processed and analyzed using Fiji/ImageJ (v1.53c). The plasma membrane and cytoplasm were defined as the peripheral region along the axonal surface and the central region of the axon, respectively. Mean fluorescence intensities of Nav1.6-EGFP and Nav1.6-tdTomato after background subtraction were obtained from these regions.

### TIRF imaging

Primary cultured hippocampal neurons at 4 DPI (21 DIV) were washed three times with 1 mL of aCSF and incubated for 15 min at 37 °C, 5% CO_2_. The coverslip was then mounted in a magnetic chamber and washed twice with 0.5 mL of aCSF before imaging.

TIRF imaging was performed as previously described (31) using an inverted microscope (IX71, Olympus) equipped with an oil-immersion objective (UPLAPO100XOHR, 100x, NA 1.50, Evident) and a 1.6* intermediate magnification lens (total magnification 160x; pixel size 100nm). Nav1.6-tdTomato near the basal membrane was excited with a 532-nm laser (OBIS 532 nm, 50 mW, Coherent) through a dichroic mirror (ZT532/640rpc, Chroma). The penetration depth of the evanescent field for the 532-nm laser was approximately 94.1 nm under our optical configuration. Emission was collected through a bandpass filter (FF01-593/40-25, Semrock) and detected with an EM-CCD camera (ImagEM, Hamamatsu). Images (512 × 512 pixels) were acquired using MetaMorph (Molecular Devices) with a 33 ms exposure time and EM gain of 1,200. Imaging was performed at room temperature.

### Analysis of single-particle diffusion

Processing of 16-bit TIFF files acquired by TIRF imaging and single-particle tracking were performed using Fiji/ImageJ (v1.53c) and the TrackMate plugin (56) following the published protocol (30,31,57). Images were preprocessed by rolling-ball background subtraction (radius = 5 pixels) and Gaussian filtering (0.5 pixels). Spot detection and trajectory reconstruction were performed using a Laplacian of Gaussian (LoG) detector and a simple linear assignment problem (LAP) tracker implemented in TrackMate, with parameters as follows: Estimated blob diameter = 4.5 pixels; Threshold = 750; Linking max distance = 3.0 pixels; Gap-closing max distance = 3.0 pixels; Gap-closing max frame gap = 1. Molecules with lifetimes exceeding two frames were included in the analysis.

Diffusion dynamics of single particles were analyzed as previously described (31). Briefly, the probability density distribution of single-particle displacements was fitted with a sum of two-dimensional diffusion equations. The parameters were optimized using the expectation– maximization (EM) algorithm. The optimal number of diffusion states was determined by Akaike information criterion (AIC). In this study, all five cells consistently indicated that a two-state model minimized the AIC. Therefore, the two-state model was adopted to describe Nav1.6 diffusion at the AIS and was uniformly applied to all diffusion analyses. The diffusion trajectories were subsequently classified into diffusion states using a hidden Markov model (HMM), yielding state occupancies.

The ensemble mean squared displacement (MSD) was calculated using the trajectories of particles in each diffusion state. The ensemble MSD curve was fitted with the circular confinement diffusion model presented as follows:

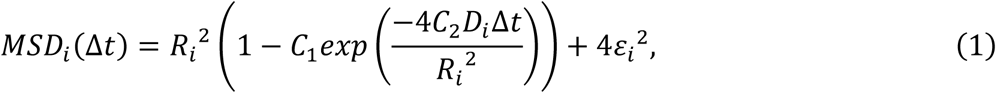

where Δ*t* represents the time lag, *D*_i_, *R*_i_, and ɛi denote the diffusion coefficient, confinement radius, and position error for each state (*i* = 1, 2), respectively. Constants *C*_1_ and *C*_2_ were set to 0.99 and 0.85, assuming a circular confinement region (58).

### Electrophysiology

For whole-cell patch-clamp recordings, CHO cells were co-transfected with either non-tagged Nav1.6 or tdTomato-tagged Nav1.6, together with Navβ1, Navβ2, and EGFP using PEI. The mass ratio of Nav1.6:Navβ1:Navβ2 is 3:1:1. Microelectrodes were made from glass capillaries (2-000-100, Drummond Scientific Company) using a puller (P-97, Sutter Instrument) and fire-polished with a Microforge MF-900 (Narishige). The microelectrodes with a resistance of 1.26– 1.75 MΩ were filled with the internal solution (105 mM CsF, 35 mM NaCl, 10 mM EGTA, 10 mM HEPES, pH 7.4 adjusted with CsOH). The bath solution was composed of 150 mM NaCl, 2 mM KCI, 1.5 mM CaCl_2_, 1 mM MgCl_2_, 10 mM glucose, and 10 mM HEPES (pH 7.4 with NaOH) (53). EGFP-positive cells were identified using an inverted fluorescence microscope (Axiovert 135, Carl Zeiss) and selected for recording at room temperature. Currents were recorded using an EPC 10 USB amplifier controlled by PatchMaster (HEKA Elektronik). Pipette offset was adjusted to zero before gigaohm-seal formation and liquid junction potential was not corrected. Capacitance transients were canceled. The speed of feedback compensation was always set to 2 μs and series resistance was compensated by 80–90% to keep the voltage error < 4.5 mV. P/N leak subtraction was not applied during recording, and baseline adjustment was applied offline during data analysis. Current recordings were initiated 3 min after break-in to minimize the effects of time-dependent shifts in channel properties. For the voltage dependence of channel activation, 50 ms depolarizing pulses were applied from −120 mV to +40 mV in 5 mV increments. For the voltage dependence of steady-state fast inactivation, 500 ms prepulses from −120 mV to +10 mV were applied in 5 mV increments, followed by a 20 ms test pulse at −10 mV.

The data were processed and analyzed using Clampfit (Molecular Devices). Peak sodium current amplitudes in each cell were normalized to the cell membrane capacitance, and the current density–voltage (*IV*) relationship are reported. Channel conductance (*G*) was calculated using the equation *G* = *I*/(*V* - *V*_rev_), where *I* is the peak sodium current amplitude, *V* is the test-pulse potential, and *V*_rev_ is the reversal potential estimated by linear extrapolation of the positive-voltage portion of the *IV* relationship for each cell. For the voltage dependence of channel activation, normalized conductances (*G*/*G*_max_) were plotted as a function of voltage and fitted with a single Boltzmann equation in the form of *G*/*G*_max_ = 1/(1 + exp((*V*_1/2,a_ -*V*)/*k*_a_)), where *G*_max_ is the maximum conductance, *V*_1/2,a_ is the half-activation voltage, and *k*_a_ is the slope factor. For the voltage dependence of steady-state fast inactivation, normalized current amplitudes (*I*/*I*_max_) were plotted as a function of voltage and fitted with a single Boltzmann equation in the form of *I*/*I*_max_ = 1/(1 + exp((*V*_1/2,i_ -*V*)/*k*_i_)), where *I*_max_ is the maximum current amplitude, *V* is the prepulse potential, *V*_1/2,i_ is the potential at which half of the sodium channels are available for activation, and *k*_i_ is the slope factor.

### Compartmental kinetic models

To quantify the kinetic constants governing asymmetric Nav1.6 turnover, the AIS was divided into proximal (0–20 μm) and distal (20–40 μm) regions based on the centroid of the Nav1.6 distribution in vitro (Fig. 3f). We then fitted the decay time courses of EGFP signals in each region using the time-evolution solutions of the following ordinary differential equations (ODEs):

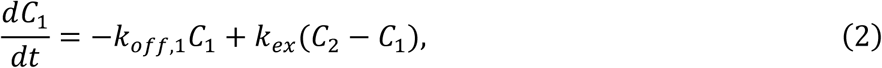

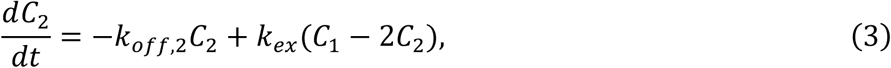

where *C*_i_ and *k*_off,i_ denote the molecular density and disappearance rate constant, respectively (*i* = 1, 2, corresponding to the proximal and distal AIS). *k*_ex_ represents the rate constant governing molecular exchange between the two compartments to account for lateral diffusion (Fig. 1i,j and Extended Data Fig. 1) (28,29). Based on previous reports showing no detectable lateral diffusion of channels between the soma and axon (28,33), we approximated *k*_ex_ at this boundary as zero in our model. In addition, because Nav1.6-EGFP or Nav1.6-tdTomato signals were extremely low or undetectable in the distal axon beyond 40 μm from the soma edge, the fluorescence intensity in this region was treated as zero. By approximating Fick’s first law with a finite-difference equation and assuming that lateral diffusion between the soma and axon can be neglected (28,33), we obtained the following relationship:

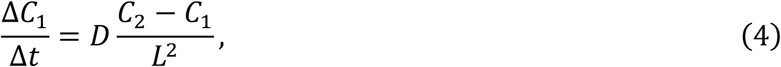

where *C*_i_ and *D* denote the molecular density and diffusion coefficient, respectively (*i* = 1, 2). Here, *L* represents the compartment length, which corresponds to the distance between adjacent equal-sized compartments, and was 19.887 μm in our analysis. By identifying this expression with the diffusion term *k*_ex_(*C*_2_ -*C*_1_) in Eq. 2, we estimated *D* from *k*_ex_ as follows:

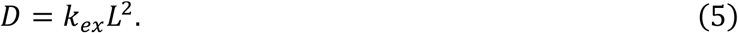

Next, with *k*_off_ and *k*_ex_ fixed at the values determined from the decay kinetics, we fitted the accumulation time courses of tdTomato signals in each region using the time-evolution solution of the following ODE:

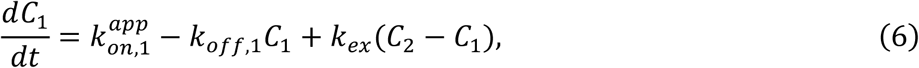

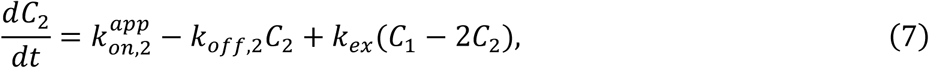

where *k*_on,i_^app^ denotes the apparent appearance rate constant (*i* = 1, 2). *k*_on,i_^app^ was normalized to *k*_on,1_^app^, yielding the relative appearance rate *k*_on,i_^rel^. The 95% confidence intervals for the estimated parameters were obtained using the bias-corrected and accelerated (BCa) bootstrap method (95% BCa CI) with 1,000 resampling iterations.

### Monte Carlo simulation

Monte Carlo simulations were performed at the single-molecule level using experimentally derived kinetic constants obtained in vitro (Fig. 3i). In the simulation, the axon was modeled as a one-dimensional axis, with 0-20 μm and 20-40 μm from the soma edge defined as the proximal and distal AIS, respectively. At each time step (Δ*t* = 0.05 d), the following three processes were iterated:

1. Molecular appearance: The number of newly appearing molecules was sampled from a Poisson distribution with mean values of 100*k*_on,1_^app^Δ*t* = 249.651 for the proximal AIS and of 100*k*_on,2_^app^Δ*t* = 187.005 for the distal AIS. The molecules were then placed randomly within each region.
2. Lateral displacement: Each molecule was moved in a random direction by a distance sampled from the one-dimensional diffusion equation with *D* = 0.0002 μm^2^ s^−1^. Following previous reports showing no detectable lateral diffusion between the soma and axon (28,33), the position at 0 μm was treated as a zero-flux boundary.
3. Molecular disappearance: The number of disappearing molecules in each region was sampled from a binomial distribution defined by the number of remaining molecules and the probability 1 -exp(-*k*_off,i_Δ*t*) for each region (*i* = 1, 2). The molecules were then removed randomly within each region.

Simulations were run up to 120 d, by which time the system had reached steady state. The net migration rate (NMR) at the proximal AIS was calculated as:

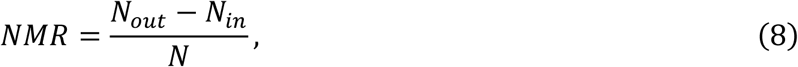

where *N*_in_ and *N*_out_ denote the number of molecules entering and leaving the proximal AIS and *N* represents the total number of molecules in the proximal AIS. The age-specific net migration contribution (ANMC) was calculated as:

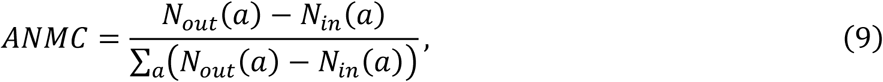

where *N*_in_(*a*) and *N*_out_(*a*) denote the number of molecules entering and leaving the proximal AIS at age *a*, respectively.

### Statistics and reproducibility

Values are expressed as mean with SD, 95% CI, or 95% BCa CI, as indicated in the figure legends. Unless otherwise noted, data are presented as mean ± SD. The number of cells or simulations used for each analysis is also provided in the figure legends. For all in vivo experiments, data from at least two separate animals were collected. Statistical significance of differences between two groups was assessed using Welch’s *t* test or paired *t* test. Dunnett’s test or Kruskal–Wallis rank-sum test was used for comparisons among more than two groups. For comparisons between proximal and distal parameters estimated from the compartmental kinetic models, significance at the 0.05 level was considered only when the 95% BCa CI for their difference did not include zero. Significance levels are \**P<0.05,* \*\**P*<0.01, and *** *P*< 0.001.

## Data availability

The data supporting the findings of this study are all available in the paper and related extended data files.

## Code availability

Custom Python code for analyzing the diffusion dynamics from single-molecule trajectories using an HMM has been deposited in GitHub (https://github.com/DaisukeYoshioka99/HMM) (57).

## Acknowledgments

We would like to acknowledge Haruo Okado (Tokyo Metropolitan Institute of Medical Science, Japan) for his invaluable advice during the initial stages of this project. We thank Meiko Kawamura and Rie Natsume (Niigata University, Japan) for insightful discussions on the design of the Nav1.6-FLEx mouse line and for technical assistance in establishing and maintaining the mouse line. We also thank Misato Yasumura, Makoto Sato (The University of Osaka, Japan), and Fumitaka Osakada (Nagoya University, Japan) for their helpful advice on AAV production. We are further grateful to Hiroshi Hibino (The University of Osaka, Japan) for his generous encouragement. This work was supported by the Center for Medical Research and Education, Graduate School of Medicine, The University of Osaka and by grants from the Japan Society for the Promotion of Science (JSPS) KAKENHI: 20K22631 and 26K18554 to D.Y.; 25K10157 to Y.Oko.; 16H06276 (AdAMS), 21K19350, 22H04922 (AdAMS), 25K02442, and 25H01334 to Y.Oka. and the Japan Foundation for Applied Enzymology to D.Y.

## Author Contributions

Y.Oka., T.K., K.S., M.A., Y.Oko., and D.Y. designed the research; D.Y., K.Y., Y.Oko., M.A., and L.H. performed the research; D.Y. carried out image data analysis and mathematical modeling; D.Y. wrote the paper.

