## Extended Data for "Asymmetric Turnover Dynamics of Nav1.6 Voltage-Gated Sodium Channels at the Axon Initial Segment"

**This PDF file includes:**

Extended Data Figures 1 to 4

### Extended Data Figures

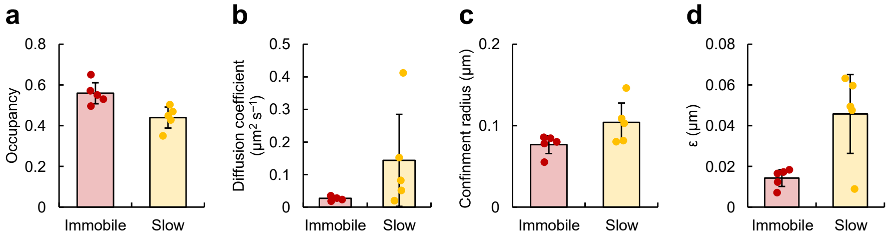


**Extended Data Fig. 1: Lateral diffusion of recombinant Nav1.6 is confined at the AIS. a–d**, Occupancies, diffusion coefficients, confinement radii, and $\varepsilon$ for immobile and slow-diffusion states of Nav1.6-tdTomato at the AIS of primary cultured hippocampal neurons at 4 DPI (21 DIV). $n$ = 5. Data are mean ± SD.

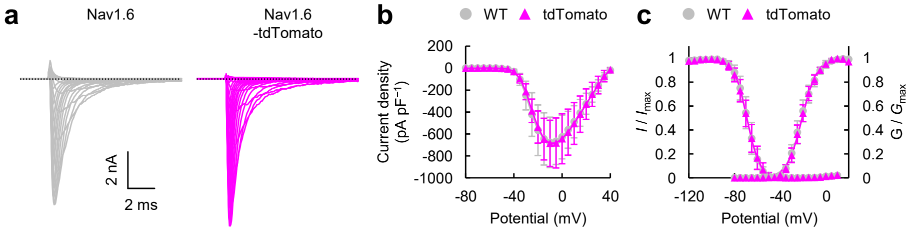


**Extended Data Fig. 2: Fluorescently tagged Nav1.6 preserves normal channel function. a**, Representative current traces of wild-type Nav1.6 and Nav1.6-tdTomato obtained by whole-cell patch-clamp recordings in CHO cells. **b**, Current densities for wild-type Nav1.6 (WT, $n = 7$) and Nav1.6-tdTomato (tdTomato, $n = 7$) at different potentials. $P > 0.05$ for all potentials (Welch's *t* test). **c**, Voltage dependence of activation and steady-state fast inactivation of wild-type Nav1.6 (WT) and Nav1.6-tdTomato (tdTomato), fitted with Boltzmann functions. $V_{1/2,\mathrm{a}}$ for WT ($n = 7$): −20.947 ± 2.529 mV and for tdTomato ($n = 7$): −20.724 ± 1.987 mV ($P = 0.868$). $k_\mathrm{a}$ for WT: 6.527 ± 0.493 mV and for tdTomato: 6.399 ± 0.220 mV ($P = 0.579$). $V_{1/2,\mathrm{i}}$ for WT ($n = 9$): −69.026 ± 3.001 mV and for tdTomato ($n = 8$): −69.163 ± 2.953 mV ($P = 0.930$). $k_\mathrm{i}$ for WT: −5.076 ± 0.369 mV and for tdTomato: −5.475 ± 0.411 mV ($P = 0.069$). *P*-values were obtained by Welch's *t* test. Data are mean ± SD.

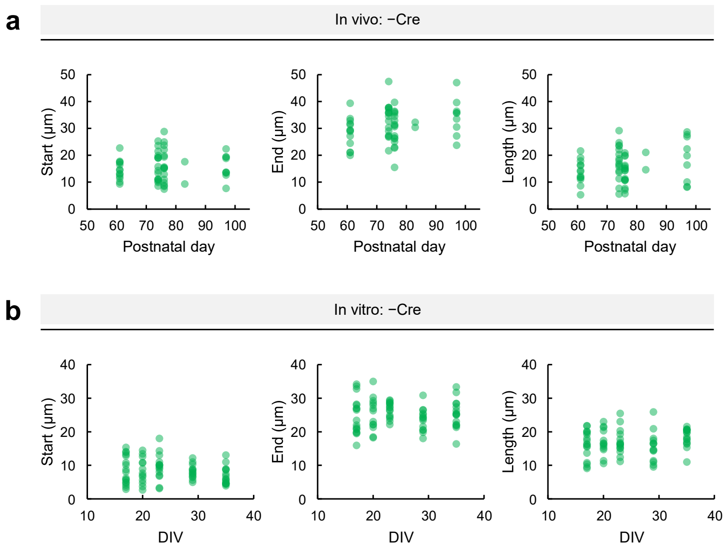


**Extended Data Fig. 3: Morphological properties of overall Nav1.6 distribution remain unchanged during in vivo and in vitro experiments. a**, Start (*P* = 0.938), end (*P* = 0.143), and length (*P* = 0.153) of the Nav1.6-EGFP distribution along the axon, measured from the edge of the nucleus, in fixed hippocampal sections without Cre expression at various postnatal days. *n* = 13 (P61), 16 (P74), 16 (P76), 2 (P83), and 9 (P97). **b**, Start (*P* = 0.270), end (*P* = 0.221), and length (*P* = 0.185) of the Nav1.6-EGFP distribution along the axon, measured from the edge of the soma, in primary cultured hippocampal neurons without Cre expression at various DIV. *n* = 17 (17 DIV), 13 (20 DIV), 15 (23 DIV), 14 (29 DIV), and 16 (35 DIV). *P*-values were obtained by Kruskal–Wallis rank-sum test.

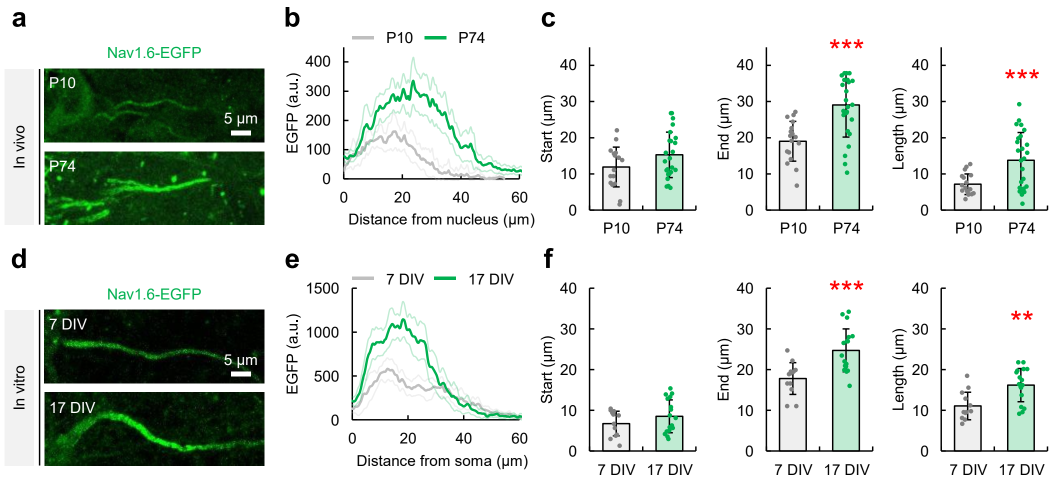


**Extended Data Fig. 4: Nav1.6 accumulates from the proximal AIS during early neuronal development. a**, Representative confocal images of neurons expressing Nav1.6-EGFP in fixed hippocampal sections without Cre expression at P10 and P74. Scale bar, 5 μm. **b**,**c**, Fluorescence intensity profiles, the start ($P = 0.079$), end (***$P = 0.000$), and length (***$P = 0.000$) of the Nav1.6-EGFP distribution along the axon, measured from the edge of the nucleus, in fixed hippocampal sections without Cre expression at P10 ($n = 17$) and P74 ($n = 25$). **d**, Representative confocal images of neurons expressing Nav1.6-EGFP in primary cultured hippocampal neurons without Cre expression at 7 and 17 DIV. Scale bar, 5 μm. **e**,**f**, Fluorescence intensity profiles, the start ($P = 0.203$), end (***$P = 0.001$), and length (**$P = 0.001$) of the Nav1.6-EGFP distribution along the axon, measured from the edge of the soma, in primary cultured hippocampal neurons without Cre expression at 7 DIV ($n = 12$) and 17 DIV ($n = 17$). *P*-values were obtained by Welch's *t* test. **b** and **e** show mean ± 95% CI; **c** and **f** show mean ± SD.
